# A Genetic Analysis of Lipid Metabolism Regulation in Han Chinese Youth in Xinjiang via Extreme Phenotypic Strategies

**DOI:** 10.1101/2025.02.18.638894

**Authors:** Jiaqing Yu, Yitong Ma

**Author notes:** **Corresponding Author** Yitong Ma, Department of Cardiology, Xinjiang Medical University Affiliated First Hospital, Wulumuqi, 830000, Xinjiang Uyghur Autonomous Region of the People’s Republic of China.

## Abstract

The Xinjiang Uyghur Autonomous Region of the People’s Republic of China has been influenced by its natural ecological environment, historical evolution, and cultural customs since ancient times. The aim of this study was to investigate new genes that contribute to lipid metabolism regulation in Han Chinese youth in the Xinjiang region through sequencing analysis. Based on the threshold value of triglycerides (TG), we selected two groups of Han Chinese university students, namely, individuals with extremely low TG levels and individuals with normal TG levels, and conducted whole-exome sequencing on samples from these individuals. Ten genes (ACTN2, DHTKD1, NLRP9, PTPRA, INPP4B, PHGDH, PYROXD2, RIN1, MYRIP, and PRSS57) were selected via bioinformatics analysis of the sequencing results. Subsequently, we can conduct further validation at the cellular and animal levels to screen for emerging genes associated with lipid metabolism.

## 1 Introduction

The Xinjiang Uyghur Autonomous Region of the People’s Republic of China has been influenced by its natural ecological environment, historical evolution, and cultural customs since ancient times. In the context of interethnic communication and biological backgrounds, the Han ethnic group exhibits an inclination towards high-fat, high-protein, and high-salt diets; these dietary inclinations are influenced by the dietary habits of the Uyghur and Kazakh ethnic groups, who predominantly consume diets that feature beef, mutton, and dairy products. This tendency is quite different from the dietary characteristics of Han populations in other provinces. The detection of novel regulatory genes that are involved in lipid metabolism and their significance in the promotion of cardiovascular health and the early treatment of dyslipidemia has consistently attracted attention in the field of internal medicine. Therefore, conducting genetic analysis studies within specific populations is more likely to reveal novel genes and new pathways. This study involved sample collection and DNA extraction from a cohort of 3,366 Han Chinese youth, followed by an analysis of the presence of lipid metabolism genes. A preliminary examination of the distribution of lipid metabolism regulatory genes in this region is reported below.

## 2 Material and methods

### 2.1 Study Subjects

The subjects who were included in this study included a total of 3,366 Han Chinese healthy young individuals who underwent health examinations at the Health Examination Center of Xinjiang Medical University from October to November 2018. Samples (including blood biochemistry, routine blood tests, surgical examination, abdominal B-ultrasound, and chest X-ray) were collected, and DNA was extracted.

### 2.2 Reagents and Instruments

#### 2.2.1 Primary Reagents Required for the Experiments Conducted in This Study and Their Respective Manufacturers

Sodium lauryl sulfate (Merck, Sigma, Inc.), ethidium bromide (Sigma, Spain), gelose (Beijing Chemical Reagents Company, China), bromophenol blue (China Shanghai Biological Engineering Co., LTD), ammonia chloride (Sinopharm Chemical Reagent Co., LTD), ammonium carbonate (Sinopharm Chemical Reagent Co., LTD), ethylenediaminetetraacetic acid (EDTA) (Sinopharm Chemical Reagent Co., LTD), chloroform (Sinopharm Chemical Reagent Co., LTD), isoamyl alcohol (Sinopharm Chemical Reagent Co., LTD), absolute ethyl alcohol (Sinopharm Chemical Reagent Co., LTD), sodium acetate (China Xinjiang Jiangweiboxin Biotechnology Co., LTD), Dynabeads® MyOne™ Streptavidin T1 (Thermo Fly, Massachusetts, USA), Agencourt SPRIselect Reagent Kit (Beckman Coulter Corporation, USA), Agencourt AMPure XP-PCR Purification Beads (Beckman Coulter Corporation, USA), Herculase®II Fusion DNA Polymerases (Agilent, California, USA), SureSelectXT Reagent Kit (Agilent, California, USA), and SSureSelectXT Human All Exon Kit V6 (Agilent, California, USA) were used.

#### 2.2.2 Relevant Consumables and Instruments Required for the Experimental Study in This Section

The following instruments and consumables were used in this study: Invitrogen Qubit 3.0 Spectrophotometer (Thermo Fisher, Massachusetts, USA), NanoDrop 2000 (Thermo Fisher, Massachusetts, USA), ABI 2720 Thermal Cycler (Thermo Fisher, Massachusetts, USA), a Covaris ME220 (American Covaris, Inc.), Agilent 2100 bioanalyzer (Agilent, California, USA), Illumina cBot Cluster Station (Inmeena, California, USA), Illumina HiSeq/NovaSeq (Inmeena, California, USA), precision electronic balance (Hko, Germany), WD-9405 type B level/upside-down rocking table (Beijing Liuyi Instrument Factory, China), DYC3ID type horizontal electrophoresis tank (Beijing Liuyi Instrument Factory, China), constant temperature metal bath instrument (Hangzhou Boi Technology Co., Ltd., China), gel imager (UVP, USA), Type U-V160 UV spectrophotometer (NanoDrop Company, USA), Eppendorf 5415 D centrifuge (E-Benz, Germany), Eppendorf 5810R PCR board centrifuge (Ebender, Germany), microbench centrifuge (Ebender, Germany), PCR instrument (MJ, USA), BECKMAN Avanti J-25 low ogenic centrifuge (Beckman Kurt, USA), −40 ℃ / −80 ℃ refrigerator (REVCO Limited, Japan), 4 ℃ refrigerator (Haier Group, China), SIM-F124 ice maker (SANYO, Japan), Magnetic Stand-96 (ABI Company), Milli-Q (Milli-Q, USA), Ultra-clean workbench (Beijing Great Wall Co., Ltd.), Eppendorff pipette (Abend, Germany), single-time centrifuge tube (50mL, 15mL, 5mL, 2mL, 1.5mL) (Ace International, Inc.), disposable gun head (0.1-1mL, 20-200uL, 0.5-20uL) (Ace International, Inc.), and vacuum collection (EDTA anticoagulation) (China Medical Device Factory).

#### 2.2.3 Preparation of Main Reagents

1. Red blood cell lysate: The chemical reagents were weighed with a precision electronic balance, and 4.9 g of ammonium carbonate, 39.9 g of ammonium chloride, and 9.9 g of EDTA were accurately measured. The weighed chemical reagents were dissolved completely in 5 liters of double-distilled water, the pH was adjusted to 8.0, and the mixture was filtered through a 0.22-µm membrane filter. The prepared reagents were sealed and stored in a refrigerator at 4 °C to prevent bacterial contamination.
2. Proteinase K solution: First, 2 g of proteinase K was weighed using a precision electronic balance and then thoroughly dissolved in 100 ml of sterilized distilled water. The prepared solution was aliquoted and stored in the dark at −20 °C.
3. Ethidium bromide working solution: First, 10 mg of ethidium bromide was weighed using a precision electronic balance and subsequently dissolved in 1 ml of sterilized distilled water. After complete dissolution, the samples were aliquoted and stored at 4 °C in a refrigerator.
4. 70% alcohol: A total of 700 milliliters of anhydrous ethanol (analytical reagent) were measured using a graduated cylinder, 300 milliliters of distilled water were added, and the volume was adjusted to 1000 milliliters. The prepared solution was subjected to repeated shaking and mixing in a 1000 mL laboratory flask.
5. l×TA Eanalytical reagent: First, a 20x TAE stock solution was prepared by accurately weighing 96.8 g of Tris hydrochloride and 14.88 g of EDTA using a precision electronic balance. The mixture was dissolved in 900 milliliters of sterilized double-distilled water and then thoroughly mixed with 114.2 milliliters of glacial acetic acid. The pH of the solution was adjusted to 8.3 using sodium hydroxide. Finally, the mixture was sealed and stored after being brought to a final volume of 1 liter in double-distilled water. Each time the working solution was utilized, 50 ml of the 100x TAE stock solution was diluted with 950 ml of double-distilled water to achieve a one-hundredfold dilution.

### 2.3 Research Ethics Review

This study was based on the major research program supported by the special fund project for central government-guided local science and technology development (ZYYD2022A01), titled “Screening for Variants of Key Genes in Dyslipidemia and Investigation of Their Regulatory Mechanisms,” specifically focusing on the key project “A Genetic Analysis of Lipid Metabolism Regulation in Han Chinese Youth in Xinjiang via Extreme Phenotypic Strategies.” The project was approved by the Ethics Committee of the First Affiliated Hospital of Xinjiang Medical University (Approval Number: 20211015-1). The Ethics Committee conducted an expedited review of the relevant materials for this study and determined that the study adheres to ethical principles. Moreover, All the participants were informed about the purpose and significance of the study, and their written consents were obtained. All the procedures were performed in accordance with the Declaration of Helsinki. Clinical trial number: not applicable.

### 2.4 Collection of Clinical Samples

This study involved physical examinations on a total of 3,366 Han Chinese undergraduate students from the first to fourth years who were enrolled at Xinjiang Medical University in 2018. Before the commencement of the health examination, we consulted relevant statistical and epidemiological experts to develop the research protocol and data collection plan, as well as to design the epidemiological data survey questionnaire and informed consent form. The information that was collected through the epidemiological data survey included, but was not limited to, personal details such as name, age, gender, height, weight, identification number, contact information, and residential address. During the implementation of the health examination, the research team conducted one-on-one surveys with participants to collect questionnaires and obtain informed consent. During the input of survey questionnaire data, two professionally trained researchers conducted cross-entry to ensure timely corrections, thereby guaranteeing the reliability of the data. Information data was managed by designated personnel, and original materials were incorporated into a database for archiving, thereby preventing information leakage.

### 2.5 Serological Testing

The subjects in the study fasted for 10–12 hours prior to the collection of blood samples, which was conducted the following morning. Blood samples were collected from the study subjects by nurses who were professionally trained at the Health Examination Center of Xinjiang Medical University. Nurses utilized EDTA anticoagulant tubes and coagulation tubes for blood collection, followed by the assessment of blood biochemistry and complete blood count. The indicators used for blood biochemistry and routine blood tests included but were not limited to red blood cell (RBC) count, white blood cell (WBC) count, platelet (PLT) count, hemoglobin (HB) concentration and other routine blood indices. Blood biochemistry mainly included TC, TG, LDL-C, HDL-C, alanine transaminase (ALT), aspartate aminotransferase (AST), and other variables.

### 2.6 Extraction of DNA from Peripheral Blood Cells

This study utilized a 5-ml whole blood genomic DNA extraction kit for the extraction of DNA, following the manufacturer’s instructions. The DNA concentration was measured after incubation at room temperature for one day. The purity and concentration of DNA were assessed using the Nanodrop2000 nucleic acid protein analyzer.

### 2.7 Second-Generation Whole-Exome Sequencing

The quality of the DNA samples was assessed using a Nanodrop 2000, and the purity of the DNA samples was evaluated through agarose gel electrophoresis. The selection and purification of DNA, fragmentation, end repair, and ligation of sequencing adapters were performed. Using the Agencourt SPRIselect reagent kit, library fragments were screened to obtain an original DNA library with fragments of appropriate length, which were subsequently subjected to PCR amplification. Hybridization of whole exome chips was performed using the SureSelectXT reagent kit. The hybrid library was then washed and purified. PCR amplification of exon DNA libraries was performed, followed by quality assessment of the DNA libraries, and sequencing was conducted using the Illumina HiSeq platform. Ultimately, the sequenced FastQ raw data were obtained. The sequencing results were analyzed using bioinformatics, and quality of the raw sequencing data was assessed. Reference sequences were analyzed by alignment with the GATK standard (https://software.broadinstitute.org/gatk/best-practices/). Detection, annotation, and classification of SNVs/InDels, followed by further filtering and prioritization of loci. This approach mainly focused on the low-frequency functional variation of genes and applied the data quality control of SNP scan typing technology provided by Shanghai Tianhao Biotech Company after sequencing.

### 2.8 Statistical Analysis

For the statistical analysis of whole-exome sequencing data and baseline characteristics of the samples, SPSS 26.0 software was used. For quantitative data, the normality of all data was initially assessed using the Shapiro-Wilk test or the Kolmogorov-Smirnov test. When the data followed a normal distribution, we employed an independent samples t test for statistical analysis. When multiple groups of data that conformed to a normal distribution were compared, one-way analysis of variance (ANOVA) was used for statistical analysis (the data are expressed as the means ± SDs). When the data did not fit a normal distribution, statistical analysis was performed using the rank-sum test. The data are expressed as the medians (interquartile ranges). For count data, we performed statistical analysis using chi-square test, and categorical data are represented using n (%). Logistic regression analysis was employed to investigate the risk factors associated with dyslipidemia. A p value of less than 0.05 was considered to indicate a statistically significant difference.

### 2.9 Trial Registration detals

Trial Registration Number: Clinical trial number: not applicable.

## 3 Results

### 3.1 Establishing a Genetic Resource Repository for Han Youth in the Xinjiang Region

Based on the previously established database of adults in the Xinjiang region, we examined the distribution of lipid profiles among adults and found significant differences in lipid levels among the Han, Uighur, and Kazakh ethnic groups: the Han population presented a relatively high HDL-C level (1.44±0.30), whereas the LDL-C level (2.19±0.60) was relatively low. However, considering that lipid levels in adults may be influenced by long-term environmental and dietary factors, confounding effects could be introduced when screening for new genes and mutation loci associated with dyslipidemia in the adult population. Therefore, conducting genetic screening studies in specific populations is more likely to reveal new genes and novel pathways (see Table 1).

**Table 1.**
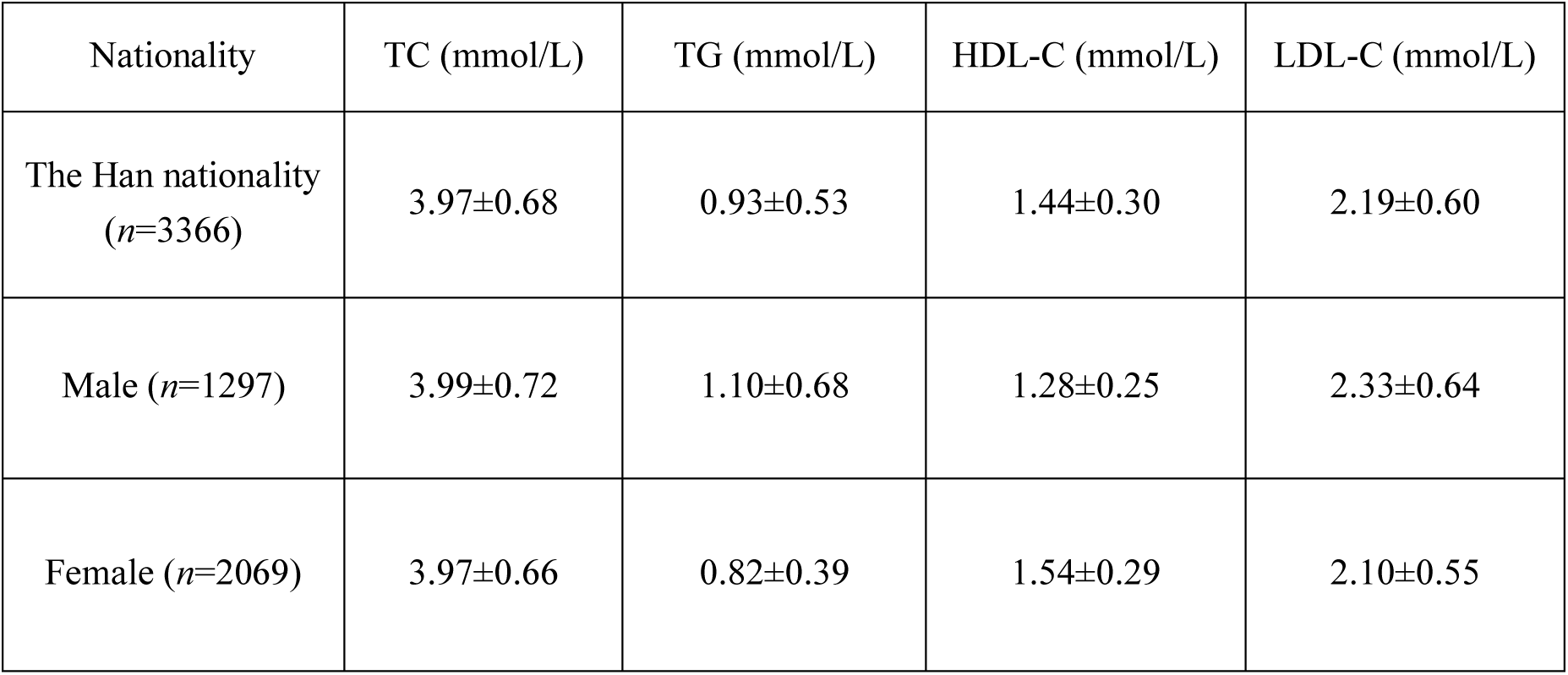
Investigation of Lipid Levels in Health Examinations of the Han Chinese Youth Population.

### 3.2 Whole-Exome Sequencing to Screen Novel Genes Involved in Lipid Metabolism Regulation

Extreme phenotype and family study strategies were adopted to rank the TG levels of Han Chinese college students. Samples from Han Chinese youth with extremely low TG levels were selected as the experimental group, and samples from Han Chinese youth with normal TG levels were used as the control group for whole-exome sequencing. After matching for gender and age, analysis of general data revealed that among Han Chinese youth, the levels of triglycerides (TG), total cholesterol (TC), low-density lipoprotein cholesterol (LDL-C), and alanine aminotransferase (ALT) in the normal TG group were significantly greater than those in the very low TG group. Conversely, the high-density lipoprotein cholesterol (HDL-C) levels in the very low TG group were significantly greater than those in the normal TG group (P < 0.05). There were no significant differences in blood glucose, AST, creatinine, and BUN levels between the two groups (see Table 2).

**Table 2.** Comparison of Baseline Data of Enrolled Han Youth after TG-level Ranking.

| Project | Normal control group<br>(*n*=102) | TG Extremely Low<br>Group (*n*=111) | Totality<br>(*n*=213) | P value |
| --- | --- | --- | --- | --- |
| Age (year) | 20.91 (1.36) | 20.76 (1.48) | 20.83 (1.42) | 0.73 |
| Sex (male,%) | 49 (48.0) | 53 (47.7) | 102 (47.9) | 0.999 |
| TG (mmol/L) | 1.43 (0.13) | 0.40 (0.04) | 0.90 (0.52) | <0.001 |
| TC (mmol/L) | 4.71 (0.38) | 3.59 (0.59) | 4.12 (0.75) | <0.001 |
| HDL-C (mmol/L) | 1.28 (0.25) | 1.56 (0.26) | 1.43 (0.29) | <0.001 |
| LDL-C (mmol/L) | 2.93 (0.23) | 1.79 (0.44) | 2.34 (0.67) | <0.001 |
| Blood glucose (mmol/L) | 4.95 (0.50) | 4.88 (0.50) | 4.91 (0.50) | 0.556 |
| Creatinine (mmol/L) | 75.11 (13.47) | 73.90 (13.44) | 74.48 (13.44) | 0.808 |
| BUN (mmol/L) | 4.40 (1.05) | 4.64 (1.12) | 4.53 (1.09) | 0.271 |
| AST (U/L) | 20.66 (11.08) | 18.69 (4.44) | 19.63 (8.35) | 0.229 |
| ALT (U/L) | 26.27 (25.80) | 15.90 (9.17) | 20.87 (19.69) | 0.001 |

After conducting a quality assessment of each sequencing dataset, we stipulated that the raw data volume for each sample should be set to ≥6G, the proportion of bases with a Q30 score should be ≥80%, and the sequencing depth of the target region should be ≥50X. Moreover, the proportion of bases with a sequencing depth of 10X or greater was set to ≥90%. The quality distribution of each base position for each sample is depicted through the original database quality distribution plot, base composition distribution plot, and sequencing error rate distribution plot. These three plots were used to visually display the sequencing quality of the R1 end of the sample to assess whether the sequencing results were evenly balanced. After the aforementioned measurements, the sequencing quality were shown to meet the standards, indicating that the sample sequencing analysis was reliable (see Figures 1–3).

**Figure 1.**
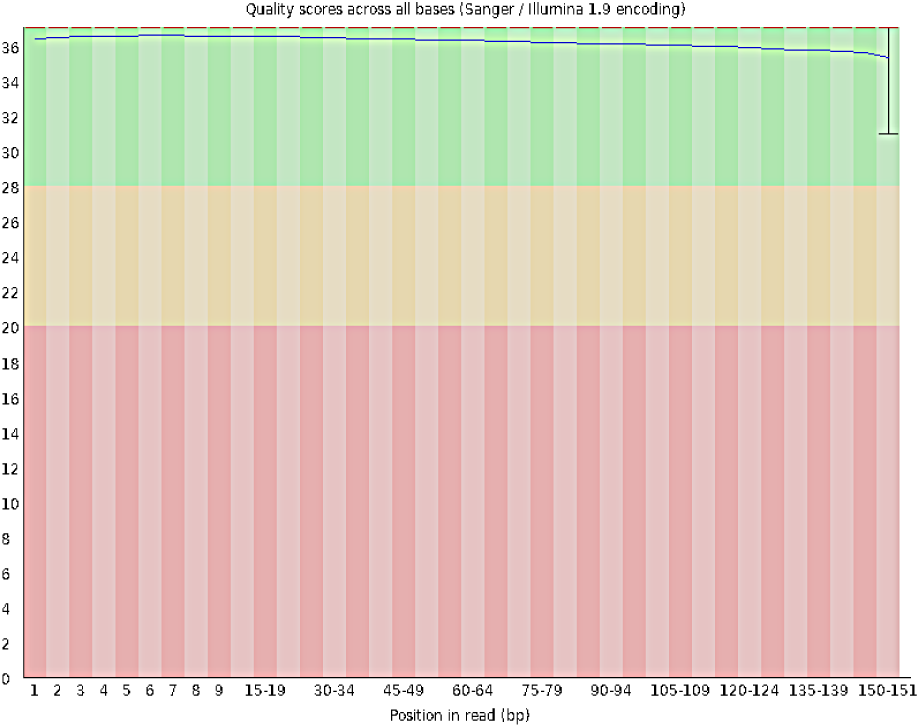
Distribution of Base Quality in the Original Data of the Han Chinese Youth Sample The horizontal axis represents the nucleotide position, and the vertical axis indicates the corresponding nucleotide quality. The background color, ranging from superior to inferior, is represented by three segments, namely, green, yellow, and red, to illustrate nucleotide quality.

**Figure 2.**
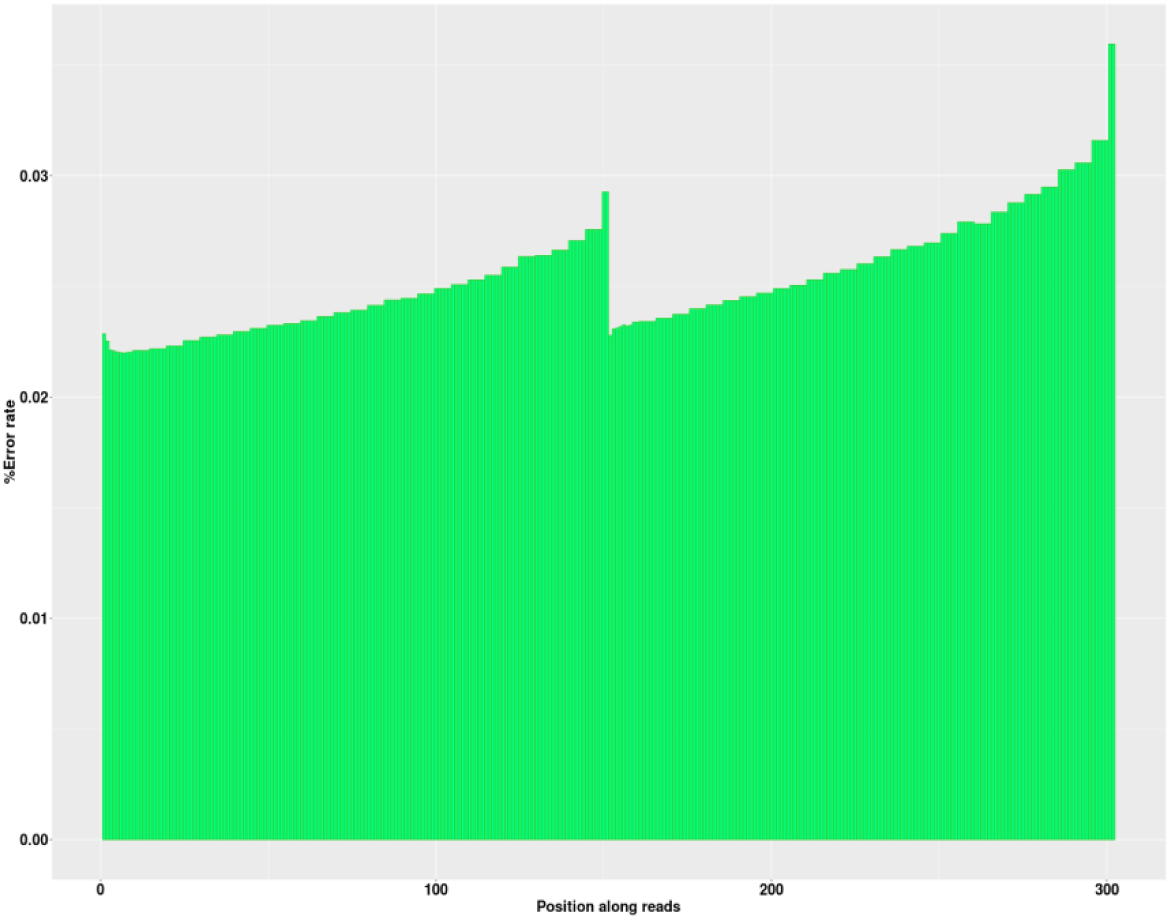
Distribution of Sequencing Error Rates in Young Han Chinese Samples The abscissa is the base position, and the ordinate indicates the base quality.

**Figure 3.**
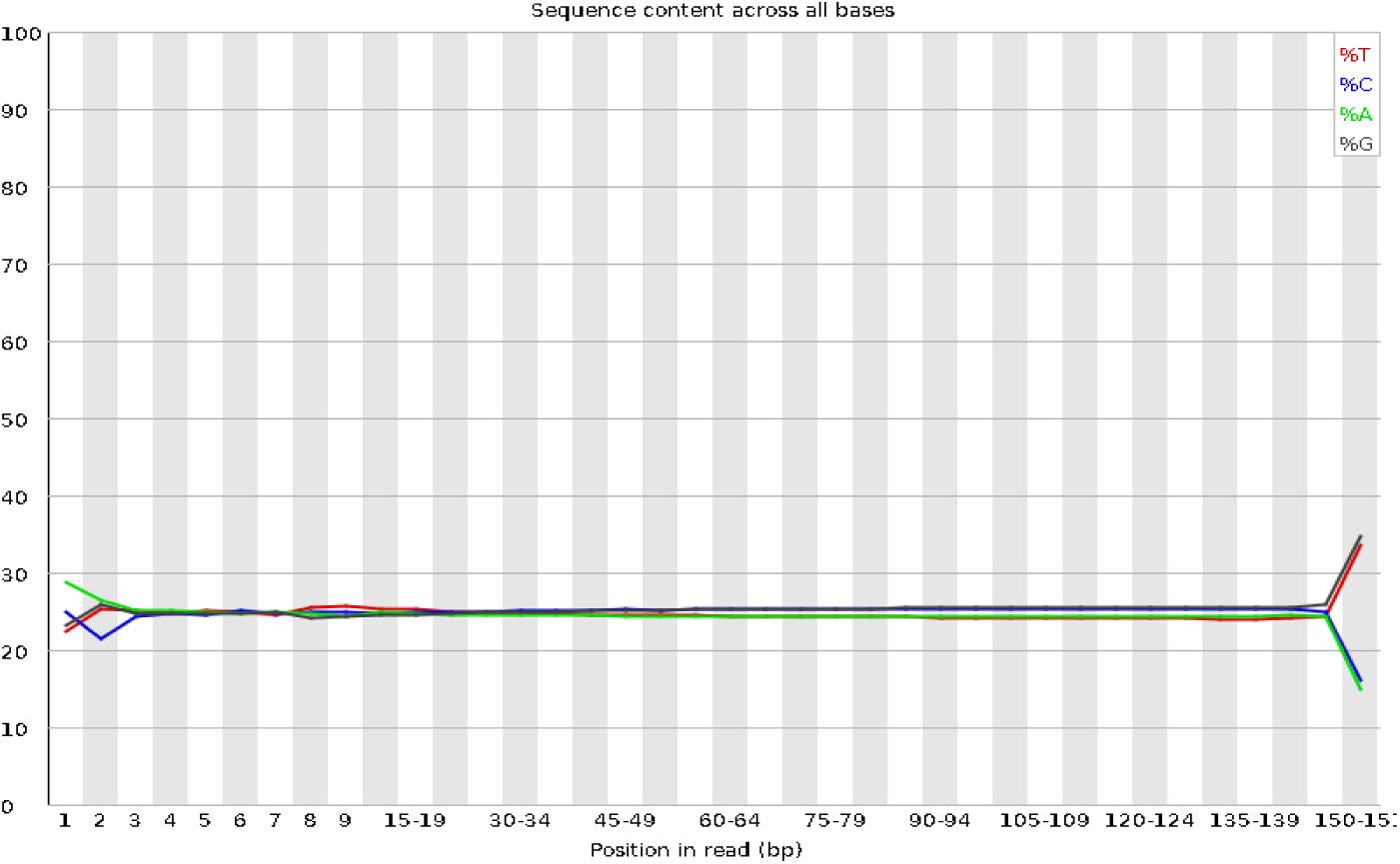
Distribution of Base Composition in Han Youth Samples The horizontal axis represents the nucleotide position, and the vertical axis indicates the proportion of individual nucleotides. Different nucleotides correspond to distinct nucleotide types.

A total of 671,127 single nucleotide variants (SNVs) were identified in the sequencing samples of Han Chinese university students. Among these, 68 mutations were located in the 5’ untranslated region (UTR) of one gene and the 3’ UTR of another gene; 231 mutations were simultaneously found in the splice region of one gene and the exon region of another noncoding RNA; additionally, 593 mutations were located in the exon region of a specific gene and the splice region of another gene. Furthermore, 765 mutations were identified in the upstream region of one gene and the downstream region of another; mutations that were located in the splice region of noncoding RNAs totaled 1,056. There were 3,344 mutations found in the downstream region of a specific gene, whereas 8,724 upstream mutations were identified in another gene. In total, 15,392 mutations were located in the 5’ UTR of one gene. The number of splice region mutations was 33,115, and the number of mutations in the exon region of noncoding RNAs totaled 15,836. Additionally, 24,516 mutations were located in the 3’ UTR, and 22,687 mutations were found in the intronic regions of noncoding RNAs. The intergenic regions exhibited 24,058 mutations. In total, 174,057 mutations were located in the exon regions, and 346,685 mutations were located in the intronic regions (Figure 4). We further classified and statistically analyzed the mutation sites based on their functional categories, which included 101 stop mutations, 937 insertion frameshift mutations, 1309 noninsertion frameshift mutations, 1949 deletion frameshift mutations, 1726 acquired stop mutations, 2609 deletions without frameshift mutations, 68799 synonymous mutations, 95839 nonsynonymous mutations, and 1381 unknown mutations (Figure 5).

**Figure 4.**
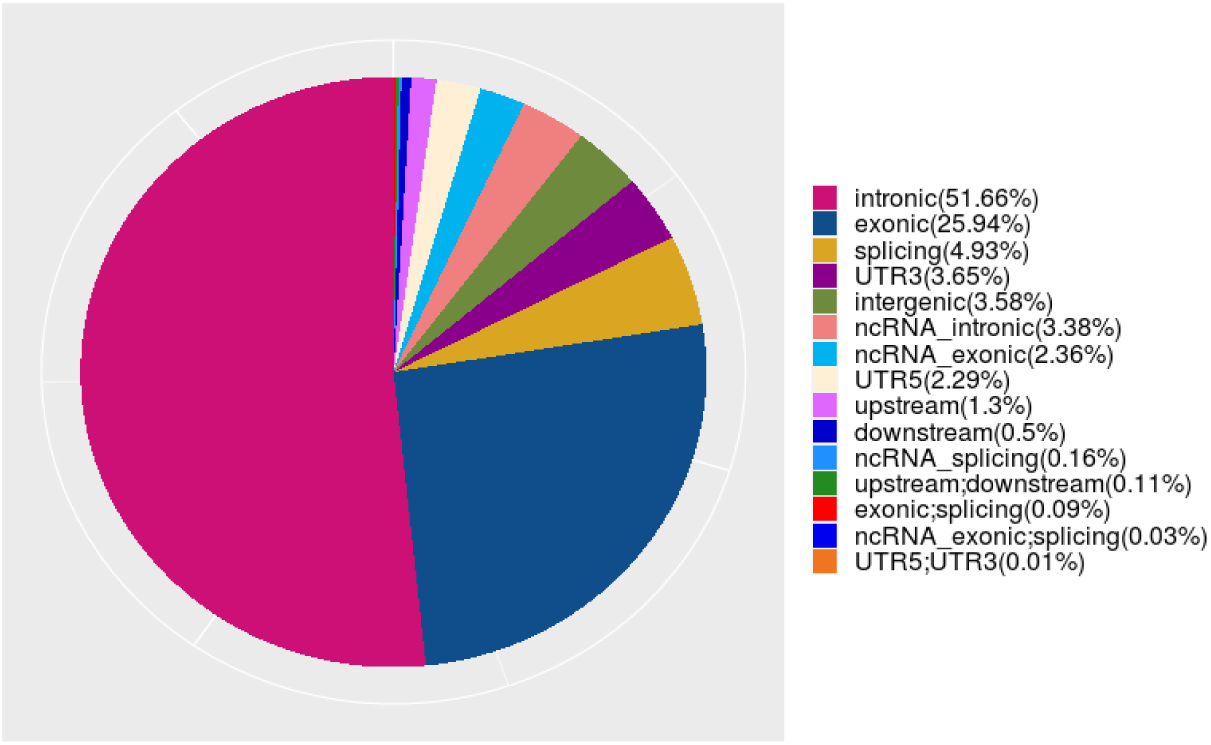
SNV Distribution of the Han Chinese Youth Sample

**Figure 5.**
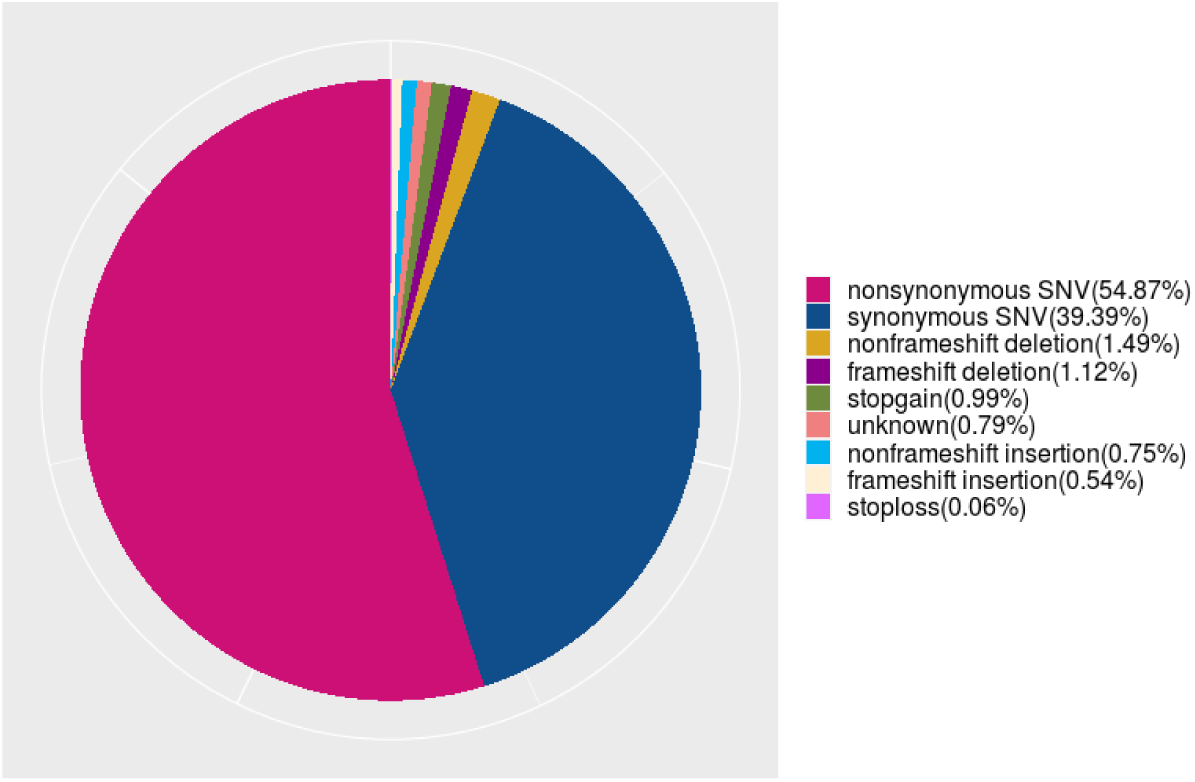
Distribution of Functional SNVs in the Sequencing Results of Han Chinese Youth

Further analysis of the insertion/deletion (InDel) mutation length distribution in sequencing samples from Han Chinese university students revealed that InDel mutations primarily occurred within the range of −5 to −1 base pairs, as illustrated in Table 3.

**Table 3.** Statistical Analysis of InDel Lengths in Han Chinese Youth.

| The number of bases inserted | Number of mutations | Percent |
| --- | --- | --- |
| 1~5 | 37442 | 36.00% |
| 6~10 | 3123 | 3.00% |
| 11~15 | 1010 | 0.97% |
| 16~20 | 617 | 0.59% |
| >20 | 852 | 0.82% |
| -5~-1 | 51494 | 49.51% |
| -10~-6 | 4468 | 4.30% |
| -15~-11 | 1953 | 1.88% |
| -20~-16 | 967 | 0.93% |
| <-20 | 2084 | 2.00% |

We continued to utilize bioinformatics to further filter the sequencing results, employing Phenolyzer software to conduct gene filtering on the mutation sites of Han Chinese university students. Through this approach, we identified candidate genes associated with disease phenotypes. The gene scores obtained by software analysis are shown in Table 4.

**Table 4.** Candidate Gene Scoring for Han Chinese Youth.

| Ranking | Gene ID | Gene name | Grade |
| --- | --- | --- | --- |
| 1 | 5566 | PRKACA | 0.9365 |
| 2 | 5594 | MAPK1 | 0.9296 |
| 3 | 7157 | TP53 | 0.9151 |
| 4 | 5578 | PRKCA | 0.9063 |
| 5 | 2033 | EP300 | 0.8978 |
| 6 | 5295 | PIK3R1 | 0.8952 |
| 7 | 805 | CALM2 | 0.8796 |
| 8 | 1387 | CREBBP | 0.8628 |
| 9 | 5568 | PRKACG | 0.8587 |
| 10 | 5291 | PIK3CB | 0.8576 |
| 11 | 5293 | PIK3CD | 0.852 |
| 12 | 5894 | RAF1 | 0.8487 |
| 13 | 4790 | NFKB1 | 0.8455 |
| 14 | 3630 | INS | 0.8224 |
| 15 | 5335 | PLCG1 | 0.8202 |
| 16 | 3845 | KRAS | 0.8155 |
| 17 | 1432 | MAPK14 | 0.7957 |
| 18 | 1956 | EGFR | 0.7951 |
| 19 | 4609 | MYC | 0.7853 |
| 20 | 5970 | RELA | 0.7753 |

We conducted a search in the HGMD database to determine whether the mutation sites associated with low-frequency functional variants of concern were already documented, which was our focus.

Additionally, we filtered for potentially impactful functional mutations in the SIFT, PolyPhenV2, MutationTaster, CADD, DANN, and dbscSNV databases, focusing on those predicted to be deleterious, located within exonic regions, or affecting splice sites. Mutations with a frequency lower than 0.001 in the 1000 Genomes database or those with a frequency less than 0.01 in self-controls were removed, and data with a frequency lower than 0.01 from the SEP6500 database were selected. Samples with a proportion of variant quality less than 20 not exceeding 50% and not classified as quality L were further filtered using the SNP Calling Quality database. Ultimately, single-nucleotide variant (SNV) loci with a homology of 1 and a mutation frequency of less than 0.005 in the Tianhao database were selected.

Ultimately, through phenotypic and genetic correlation analyses, we identified genes that were potentially associated with cholesterol metabolism (primarily affecting triglycerides). Based on the statistical results, we selected the top ten genes ranked by their effects (see Table 5).

**Table 5.** Top 10 Candidate Genes in Han Chinese Youth A: Wild-type samples; B: Heterozygous mutant samples; C: Homozygous mutant samples.

| Gene | SNV number | Number of mutation samples | Total mutation (A\|B\|C) | Extreme low group (A\|B\|C) | Contrast group (A\|B\|C) | Mutation rate | *P* |
| --- | --- | --- | --- | --- | --- | --- | --- |
| ACTN2 | 10 | 10 | 199 \| 9 \| 1 | 98 \| 9 \| 1 | 101 \| 0 \| 0 | 10 | 0.00228 |
| DHTKD1 | 12 | 12 | 197 \| 12 \| 0 | 97 \| 11 \| 0 | 100 \| 1 \| 0 | 10.287 | 0.00532 |
| NLRP9 | 10 | 13 | 196 \| 12 \| 1 | 97 \| 11 \| 0 | 99 \| 1 \| 1 | 5.143 | 0.00532 |
| PTPRA | 7 | 12 | 197 \| 12 \| 0 | 97 \| 11 \| 0 | 100 \| 1 \| 0 | 10.287 | 0.00532 |
| INPP4B | 4 | 8 | 201 \| 8 \| 0 | 100 \| 8 \| 0 | 101 \| 0 \| 0 | 8 | 0.00702 |
| PHGDH | 3 | 8 | 201 \| 8 \| 0 | 100 \| 8 \| 0 | 101 \| 0 \| 0 | 8 | 0.00702 |
| PYROXD2 | 6 | 8 | 201 \| 8 \| 0 | 100 \| 8 \| 0 | 101 \| 0 \| 0 | 8 | 0.00702 |
| RIN1 | 11 | 12 | 197 \| 11 \| 1 | 97 \| 10 \| 1 | 100 \| 1 \| 0 | 10.287 | 0.00708 |
| MYRIP | 10 | 11 | 198 \| 11 \| 0 | 98 \| 10 \| 0 | 100 \| 1 \| 0 | 9.351 | 0.01018 |
| PRSS57 | 8 | 11 | 198 \| 11 \| 0 | 98 \| 10 \| 0 | 100 \| 1 \| 0 | 9.351 | 0.01018 |

## 4 Discussion

The relationship between dyslipidemia and cardiovascular diseases has been substantiated in epidemiological and evidence-based medical studies. To improve the occurrence and progression of cardiovascular diseases, early prevention of dyslipidemia is considered a significant research direction in the field of cardiology. Although dyslipidemia is commonly associated with an increased risk of cardiovascular diseases, it should be noted that isolated dyslipidemia does not immediately manifest clinical features of cardiovascular conditions. Rather, the effects accumulate gradually over time with prolonged exposure, ultimately leading to the onset of cardiovascular events such as acute coronary syndrome and stroke. With the aim of facilitating the early detection, early prevention, and early treatment of dyslipidemia and metabolic disorders, extensive research has been conducted in the field of cardiovascular internal medicine. Two hundred years ago, Michel Eugène Chevreul extracted C27H46O from human gallstones and named it cholesterol. In 1910, foreign scientists discovered that the cholesterol content in atherosclerotic plaques of the human aorta was 25 times greater than normal values. The academic community subsequently initiated in-depth research into the relationship between atherosclerosis and increased dietary cholesterol. This potential relationship was corroborated by Nikolaj Anitschkow in his studies on atherosclerosis induced by cholesterol feeding in rabbits (1). Since the beginning of the 21st century, significant progress has been made in research on cholesterol metabolism. At present, it is believed that cholesterol exists in the body, and its homeostasis is closely monitored. The dynamic balance of exogenous intake (food acquisition), biosynthesis, output, and esterification of cholesterol is reflected in the cellular cholesterol content. Among these four processes, the first two— food acquisition and biosynthesis—are the primary sources of cholesterol, primarily facilitated by the Niemann-Pick C1-like 1 (NPC1L1) protein in the epithelial cells of the small intestine. NPC1L1 functions as a cholesterol importer and is involved in the intestinal absorption of cholesterol. NPC1L1 is also involved in the regulation of the conversion of dietary cholesterol intake (2). After the depletion of intracellular cholesterol, the NPC1L1 protein associates with the outer Flotillin-1 and Flotillin-2 proteins with the assistance of the Numb protein and Clathrin/AP2 proteins, while Myosin Vb also binds to the NPC1L1 protein. This process involves the participation and assistance of the LIM domain and actin-binding protein 1 (LIMA1) (3). Cholesterol can also be transported into the cell by cytoplasmic microfilaments to mitigate cholesterol depletion, which is a process that relies on the involvement of NPC1L1 (4–8). When cholesterol is present in excessive levels, the apolipoprotein A-I (APOA-I) produced by the liver, intestines, and pancreas can bind to cholesterol in the body to form high-density lipoprotein (HDL) (9). Acyl-CoA: Cholesterol acyltransferase (ACAT) has also been found to esterify excess cholesterol into cholesteryl esters for storage in cellular lipid droplets. The detection of VLDL and LDL in the blood is also associated with ACAT. The liver and intestines are also capable of clearing HDL, which is returned via the circulatory system (10). This series of processes is considered to represent the dynamic equilibrium of cholesterol. Very low-density lipoproteins (VLDL) in the bloodstream have been identified as a form of cholesterol that is endogenously synthesized and exogenously absorbed by the liver (11), subsequently transforming into low-density lipoproteins (LDL) and triglycerides (TG) (12) in circulation. Dyslipidemia is also a critical factor that contributes to the development of atherosclerotic diseases.

Cholesterol is located within cell membranes and is an amphipathic molecule that is characterized by its hydrophobic properties. Cellular fluidity, permeability, and polarity can be modulated following the interaction of cholesterol with adjacent lipid species, and cholesterol can associate with various transmembrane proteins to adjust protein conformation. The regulation of cholesterol transport and subcellular distribution can increase efficiency under the influence of sterol transport proteins (13). The Hedgehog signaling pathway has been studied and is believed to be regulated by cholesterol, which is considered an important biomolecule necessary for the structural integrity of cell membranes, cellular signaling, and precursors for steroid hormones and bile acids. Cholesterol is thought to covalently modify Hedgehog, thereby achieving regulatory effects (14–16). We believe that interventions can be administered to target process and influence cholesterol balance and that cholesterol synthesis, which is regulated by multiple genes, can be modulated.

Sterol regulatory element-binding protein 2 (SREBP2) is of paramount importance, as it plays a crucial role in the transcriptional regulation of enzymes associated with the cholesterol synthesis pathway. The enzymes associated with the cholesterol synthesis pathway include more than twenty distinct enzymes, which is why it is often considered a central process (17). The SREBP cleavage-activating protein (SCAP) in the endoplasmic reticulum interacts with SREBP2 (18). To address the depletion of cholesterol in the endoplasmic reticulum, SCAP translocates to the Golgi apparatus after binding to coatomer II (COPII) (19). The proteases S1P and S2P cleave SREBP2, releasing the N-terminal domain, which subsequently binds to the sterol regulatory element (SRE) and activates transcription-related genes (18).

On the other hand, 3-hydroxy-3-methylglutaryl-coenzyme A reductase (HMGCR) is a rate-limiting enzyme in cholesterol synthesis. As the catalyst for the first rate-limiting step in the cholesterol biosynthetic pathway, HMGCR has been shown in previous studies to be regulated by SREBP2, which maintains cholesterol homeostasis. This regulation is often achieved through the specific binding of SREBP2 to the SRE-1 element of the HMGCR gene during the process (20). The ubiquitination of cholesterol is of paramount importance, as ubiquitinated cholesterol is subject to degradation by the proteasome (21). Research suggests that the insulin-induced gene (INSIG) protein interacts with cholesterol HMGCR in the process of ubiquitination. When cholesterol levels are elevated, elevated cholesterol levels form a SCAP-SREBP2-INSIG complex with SREBP cleavage-activating protein, which subsequently interacts with HMGCR, leading to ubiquitination and degradation within the proteasome. When intracellular cholesterol is depleted, INSIG no longer binds to increase cholesterol synthesis (22–25). Another mechanism has also been reported: ATP-binding cassette subfamily G member 5 and member 8 (ABCG) transporters can secrete cholesterol into the extracellular space upon binding ATP (26–28). When cholesterol levels are excessive, ATP-binding cassette subfamily A member 1 (ABCA1) can bind ATP to facilitate the efflux of free cholesterol from the cell, subsequently associating with APOA-I to form high-density lipoprotein (HDL). The HDL formed through this active/passive pathway enters the circulatory system (29, 30).

Currently, the lipid-lowering agents that are extensively studied and utilized in clinical practice include statins, ezetimibe, and PCSK9 inhibitors. The pharmacological effects of these drugs are aimed at reducing blood lipids through three pathways: 1) Statins inhibit the endogenous synthesis of cholesterol by targeting HMGCR; 2) ezetimibe, which is a targeted drug, inhibits intestinal NPC1L1, thereby reducing the absorption of cholesterol once it enters the intestine; and 3) PCSK9 inhibitors, which function as antibodies against the PCSK9 protein, bind to PCSK9 and inhibit its interaction with LDLR, leading to the degradation of LDLR in lysosomes. Currently, statins are utilized as the primary pharmacological intervention in clinical practice. However, a subset of patients who exhibit statin resistance, characterized by dyslipidemia despite adherence to the target dosage, has been identified. For these individuals, combination therapy involving ezetimibe and PCSK9 inhibitors may be considered.

Blood flow can be improved following the administration of lipid-lowering medications, and this intervention may significantly reduce the risk of thrombus formation. It has been reported that statins combined with ezetimibe can increase blood lipid reduction levels (29) with guaranteed safety. Patients with cardiovascular diseases can benefit from the combination of statins and PCSK9 inhibitors; the risk of developing cardiovascular diseases and strokes is reduced by 15%, and lipid levels are decreased by 60% (30). Therefore, the combination of statins and PCSK9 inhibitors in the treatment of cardiovascular diseases can slow disease progression and contribute to a reduction in the incidence of cardiovascular events. However, there are certain limitations associated with these medications. For example, when ezetimibe is used as a monotherapy, its efficacy in lowering cholesterol levels is suboptimal, and it may lead to gastrointestinal reactions such as diarrhea, nausea, and abdominal pain, as well as potential liver damage and myalgia. The use of statins can result in muscle pain, cramps, and, in severe cases, rhabdomyolysis, as well as possible hepatic impairment. Additionally, the high cost of PCSK9 inhibitors currently precludes their widespread adoption, which may lead patients to discontinue treatment prematurely, thereby interrupting the therapeutic process. The medical community is very interested in discovering new lipid-lowering agents; however, current pharmaceutical research related to genes that are involved in cholesterol metabolism has encountered numerous challenges and setbacks. Cholesteryl ester transfer protein (CETP) inhibitors, which are an emerging class of lipid-lowering agents, have been the subject of extensive research. CETP is produced by the liver as a glycoprotein and can preferentially bind to HDL. This protein subsequently transforms into LDL and VLDL after exchanging chylomicrons with APOB. Currently, five types of inhibitors have been developed: TorcetRapib, DalcetRapib, EvacRapib, Anacetrapib, and ObicetRapib; research has shown that Torcetrapib is associated with an increased mortality rate in cardiovascular populations (31). Conversely, Dalcetrapib and Evacetrapib have been confirmed in studies to have no significant effect (32, 33). The latter two drugs are still undergoing clinical trials. In summary, to investigate new lipid-lowering medications, it is essential to continue the search for novel genes that are associated with cholesterol homeostasis and to explore their functional mechanisms.

At the beginning of the new century, the rapid iteration and upgrading of modern sequencing technologies have facilitated the widespread application of bioinformatics techniques and large-scale genomic sequencing analyses for the identification of gene mutations within high-throughput platforms. For example, the Global Lipids Genetics Consortium (GLGC) has identified 157 mutation loci in its lipid research, including a substantial sample size of 187,000 human subjects (34). GWAS studies, as the predominant research approach for identifying mutation sites associated with risk factors, are capable of elucidating the correlation between phenotypes and genetic mutations. GWASs have previously identified hundreds of variant loci that are associated with cardiovascular diseases and lipid levels. Research on lipid-related genes across different ethnic populations is also underway. In the GWAS conducted by Themistocle with a sample of 312,571 individuals to investigate novel mutation loci that are associated with lipid profiles, a total of 118 new mutation loci related to lipid metabolism were identified. However, certain variants exhibited inconsistencies across different ethnic groups, such as variations found in European populations compared with those in nonEuropean populations (35). However, while GWASs can contribute significantly to the exploration of drug targets, they still have certain limitations. Due to limitations imposed by traditional analytical methods, the effects of novel mutation loci that are identified in GWASs are relatively weak, which fails to fully account for the variability of hereditary traits and accurately pinpoint the key genes that influence a specific phenotype. Therefore, although a GWAS can provide new insights into drug targets, it is not able to accurately identify the key genes that affect a certain phenotype to completely justify the causal correlation of the phenotype with the gene. Therefore, additional biological functional experiments are necessary for relevant validation.

Historically, the primary approach has involved comparing sample sequencing between populations with hyperlipidemia and control groups, thereby identifying potential genetic mutations associated with hyperlipidemia for further investigation. However, the population affected by hyperlipidemia primarily consists of middle-aged and elderly individuals, with hyperlipidemia resulting from a combination of environmental and genetic factors being most prevalent in this population.

However, individual lifestyle habits can render environmental factors difficult to control, making it impossible to fully regulate the duration of exposure to risk factors. Consequently, this inability to exclude such confounding variables during the analysis of sequencing results may introduce a degree of bias in the evaluation of genetic mutations that affect lipid levels. Due to the relatively short duration of environmental and dietary influences on lipid levels in the youth population, most of whom do not engage in habits such as smoking or alcohol consumption and lack other adverse underlying health conditions, we posit that new genes may be more readily identified during genetic screening within this population.

Therefore, this study selected young university students as the research subjects, which ensures that their phenotypes primarily arise due to genetic factors, thereby eliminating the confounding effects of environmental factors. This also represents an innovative aspect of the research.

## 5. Conclusions

This study employed a combined strategy of extreme phenotype and family-based research, selecting Han Chinese university students from the Xinjiang region. The participants were ranked according to their triglyceride (TG) levels, from highest to lowest. The study focused on the Han Chinese subgroup with extremely low TG levels and a normal TG group (Han Chinese: control group: 102 individuals, extremely low TG group: 112 individuals, TG ≤ 0.45 mmol/L). This study focused on low-frequency functional mutations, which occur at a very low probability within the population. Based on the threshold value of triglycerides (TG), we selected two groups of Han Chinese university students, individuals with extremely low TG levels and those with normal TG levels, whole-exome sequencing was performed on samples from these individuals. Ten genes (ACTN2, DHTKD1, NLRP9, PTPRA, INPP4B, PHGDH, PYROXD2, RIN1, MYRIP, and PRSS57) were selected via bioinformatics analysis of the sequencing results. We sequenced approximately one hundred samples from the Han Chinese population. The results of low-frequency functional new gene screening among Han Chinese university students indicated that, due to historical long-term integration and exchange, the genetic background and the genes influencing lipid metabolism in the Han ethnic group are relatively complex. Due to the collective influence of multiple genes, the effect of each individual gene is limited. Therefore, the identification of known classical genes that are associated with lipid metabolism via this extreme phenotype strategy is more time-consuming and poses greater challenges. Therefore, screening for known classical genes that are associated with lipid metabolism via this extreme phenotype strategy would be more time-consuming and challenging. Subsequently, we can conduct further validation at the cellular and animal levels to screen for emerging genes associated with lipid metabolism.

## List of Abbreviations

TG: triglycerides
RBC: red blood cell
WBC: white blood cell
PLT: platelet
Hb: hemoglobin
AST: aspartate aminotransferase
ALT: alanine transaminase
TC: total cholesterol
LDL-C: low-density lipoprotein cholesterol
HDL-C: high-density lipoprotein cholesterol
ACAT: Acyl-CoA: cholesterol acyltransferase
VLDL: very low-density lipoproteins
LDL: low-density lipoproteins
SREBP2: sterol regulatory element-binding protein 2
SCAP: SREBP cleavage-activating protein
SRE: sterol regulatory element
COPII: coatomer II
SRE: sterol regulatory element
HMGCR: 3-hydroxy-3-methylglutaryl-coenzyme A reductase
INSIG: insulin-induced gene
ABCG: ATP-binding cassette subfamily G
ABCA1: ATP-binding cassette subfamily A member 1
CETP: cholesteryl ester transfer protein
GLGC: Global Lipids Genetics Consortium.

## Declarations

### Human Ethics and Consent to Participate declarations

#### The project was approved by the Ethics

Committee of the First Affiliated Hospital of Xinjiang Medical University (Approval Number: 20211015-1). The Ethics Committee conducted an expedited review of the relevant materials for this study and determined that the study adheres to ethical principles. Moreover, All the participants were informed about the purpose and significance of the study, and their written consents were obtained. All the procedures were performed in accordance with the Declaration of Helsinki.

## Consent for Publication

All authors consent to publication

## Availability of data and material

All data and material supporting this work will be available upon reasonable request from the corresponding author.

## Authors’ Contributions

Q.-J.Y. conceived the ideas, designed the experimental procedures, analyzed the data, performed the majority of the experiments, prepared the manuscript and edited the manuscript. Q.-J.Y. conceived the experiment and reviewed the manuscript. T.-Y.M. critically reviewed the manuscript.

## Acknowledgments

The authors would like to thank AJE (https://www.aje.cn/) for the English language review. This work was supported by the special fund project for central government-guided local science and technology development (ZYYD2022A01).

